# Detection of Infectious Murine Norovirus on Fresh Produce through a Novel CRISPR/Cas13-based Sensitive Assay

**DOI:** 10.1101/2025.04.04.647263

**Authors:** Axel Ossio, Angel Merino-Mascorro, Juan S. Leon, Norma Heredia, Santos Garcia

**Affiliations:** Universidad Autónoma de Nuevo León, Facultad de Ciencias Biológicas. Laboratorio de Bioquímica y Genética de Microorganismos, San Nicolás de los Garza, N.L.,México 066455; Hubert Department of Global Health, Rollins School of Public Health, Emory University

**Keywords:** Norovirus, CRISPR-Cas13a, foodborne, infectivity, lettuce, blueberries

## Abstract

Current standard food detection methods do not distinguish between infectious and non-infectious norovirus leading to uncertainty in the interpretation and management of a norovirus positive food sample. These methods also require expensive RT-qPCR based equipment. In contrast, CRISPR-based, compared to RT-qPCR based, detection methods yield similar sensitivity and specificity and are generally less expensive. The aim of this study was to detect norovirus with an intact capsid, a proxy for infectivity, through a CRISPR-Cas13a based detection method in conjunction with acapsid integrity assay. Our CRISPR method detected murine norovirus (MNV-1), with an intact capsid, at a limit of detection of 2.59 log10 gc/ 25 g (5 gc/ rx). This method did not cross-react with other targets (synthetic hepatitis A virus; human norovirus GI, GII; rotavirus). Compared with RT-qPCR, this CRISPR based method showed an increased sensitivity when detecting low copy numbers of RNase-pretreated MNV-1 in lettuce and blueberries samples. This is the first report describing a CRISPR-based detection of potentially infectious viruses in food samples.

## 1. Introduction

Human norovirus is the most common cause of acute gastroenteritis worldwide, with an estimated 685 million cases annually (Wang et al., 2023). Noroviruses are non-enveloped, single-stranded RNA viruses that infect a wide range of mammalian host species, including humans (Randazzo et al., 2018). Based on phylogenetic clustering of the complete VP1 amino acid sequence, noroviruses are classified into ten genogroups, GI-GX (GI, GII, and GIV can cause human infections), and 48 genotypes (Chhabra et al., 2019; Khan and Alam, 2021).

Norovirus is considered an important foodborne pathogen in outbreaks associated with produce (Bosch et al., 2018) including fruits (e.g., berries) and vegetables (e.g., leafy greens), which are often consumed raw or minimally processed (Yeargin and Gibson, 2019). Contamination of norovirus, due to poor hygiene, may occur at any point of the “farm to fork” supply chain, including handling during harvest, postharvest and food preparation by handlers (Sobolik et al., 2021). Norovirus is stable under different conditions used in the food industry such as high and low temperatures, extreme pH, pressure conditions and to many chemical disinfectants and this stability also contributes to its transmission (Cook et al., 2016). Further, the ID50 of only 18 infectious viral particles requires sensitive detection methods of low viral copies in food samples. Additionally, methods to discriminate between infectious and non-infectious viral particles are needed to determine the true risk of a food with a functional replication-able norovirus versus environmentally persistent norovirus nucleic acids with a damaged or absent capsid (Knight et al., 2013; Sobolik et al., 2021).

Among viral detection methods in environmental and food samples, RT-qPCR is the gold standard. However, because RT-qPCR detects only genetic material, it does not discriminate between norovirus genetic material in a functional capsid, a non-functional capsid, or from free environmentally persistent RNA (Chen et al., 2020). Human norovirus cell culture systems can detect infectious norovirus but lack sensitivity and cannot be applied to all norovirus strains. Thus, several groups have developed indirect detection methods of infectivity, including molecular approaches to detect intact viral capsids, a proxy of infectivity (Moore et al., 2017). Some of these approaches includes ligand binding-based assays using synthetic human norovirus histo-blood group antigen (HBGA) or porcine gastric mucin, and capsid and genome integrity assays such as RT-qPCR preceded by RNase and/or proteinase and the use of photo activatable intercalating agents like ethidium monoazide and propidium monoazide (Liu and Moore, 2020; Manuel et al., 2018; Moore et al., 2017).

In addition to RT-qPCR, other sensitive detection methods of viral genetic material include isothermal amplification methods (RPA, LAMP), whole genome or amplicon-based sequencing methods and droplet digital PCR (ddPCR) (Chin et al., 2022; Raymond et al., 2022; Suther et al., 2022). Although these technologies offer high specificity and sensitivity, like RT-qPCR, they cannot discriminate between infectious and non-infectious viruses and are equally expensive and time consuming. For example, all these technologies require expensive and sophisticated laboratory equipment together with specialized and trained personnel in their use.

The Clustered Regularly Interspaced Short Palindromic Repeats, when coupled with Cas enzymes, offers an alternative to traditional RT-qPCR methods in detecting pathogens (Zhu et al., 2020). These CRISPR Cas systems are not only comparable in sensitivity and specificity to RT-qPCR but also boast advantages such as reduced assay times, lowered costs, and the elimination of the need for costly RT-qPCR equipment (Palaz et al., 2021; M. Wang et al., 2020; X. Wang et al., 2020). Moreover, they can be integrated into laboratories currently performing RT-qPCR assays. The application of CRISPR with the Cas13a enzyme, for example, has demonstrated the capability to identify as few as 5 genomic copies of norovirus GII.4 from stool samples within two hours at a temperature of 39°C (Duan et al., 2022). On foods, there have been CRISPR-Cas-based assays to detect *E. coli* O157:H7, *Listeria monocytogenes, Salmonella* Typhimurium, *Staphylococcus aureus*, and *Vibrio parahaemolyticus* (Hadi et al., 2023; Lu et al., 2022), and recently, for human norovirus and rotavirus on lettuce and strawberries (Le et al., 2025). Despite these advances, the detection of viral RNA by CRISPR does not provide data on viral infectivity, thereby necessitating the integration of an RNase pretreatment step, as proposed in this study.

There is a need for methods that are as sensitive and specific as RT-qPCR, are less expensive and laborious, and can distinguish non-infectious norovirus (Liu and Moore, 2020; Raymond et al., 2021). Such methods would enhance the food industry’s access to viral detection tools and improve their source tracking of infectious norovirus and interpretation and management of a food sample with non-infectious norovirus. Given this need, the aim of this study was to detect norovirus with an intact capsid, a proxy for infectivity, through a CRISPR-Cas13a based detection method in conjunction with a capsid integrity assay. We evaluated this system on murine norovirus (MNV-1), as a human norovirus surrogate, on fresh produce, including blueberries and lettuce (Ossio, et al., 2024).

## 2. Materials & methods

### Viral stock

Murine norovirus CW1 strain (MNV-1, ATCC^®^ VR-1937™) was used as a human norovirus surrogate. MNV-1 was propagated in RAW 264.7 cells (ATCC^®^ TIB-71™) maintained in Dulbecco’s Modified Eagle’s Medium” (DMEM) (Sigma-Aldrich^®^ St. Louis, MO) supplemented with 10% FBS, penicillin (100 U/ml) and streptomycin (100 µg/ml) (Sigma-Aldrich^®^) at 37 °C with 5% CO_2_. Stocks were obtained after consecutive rounds of virus infection in confluent cells (75-cm^2^ culture flasks). Cells were scraped and the suspension was subjected to three freeze-thaw cycles followed by clarification using low-speed centrifugation of 1 460 × g for 15 min. Further, cells were adjusted and infected at a multiplicity of infection (MOI) of 0.1. The viral aliquots were stored at -80 °C. Viral titrations were performed following the RT-qPCR protocol.

### Sample collection and inoculation

Fresh iceberg type lettuce (*Lactuca sativa var. capitata*) and the common North American blueberry (*Vaccinium corymbosum*) were bought at local food stores in Northern Mexico. These food matrices were selected as representative fruits (e.g., berries) and vegetables (e.g., leafy greens) associated with norovirus outbreaks (Cook et al., 2019; Sobolik et al., 2021; Torok et al., 2019; Yang and Scharff, 2024). Both food matrices were independently subdivided in 25 g samples and placed in Whirl-Pak^®^ bags (VWR™, WI, USA). Any adhered soil was removed by gentle scrubbing. Six ten-fold diluted doses (5.06 × 10^5^ to 5.06 genomic copies/ reaction [gc/ rx] in 300 μl of PBS) of stock murine norovirus inoculum (5.06 x 10^9^ gc/ rx) were spot inoculated on replicate 25g produce samples and allowed to air dry for 1 h. For both food matrices, a no-virus inoculum control of 270 μl PBS with was included as negative control. For each food matrix and each dilution, three independent experiments with three replicates each were performed.

### Virus elution and concentration

For elution and concentration from each food sample, the ISO 15216-1:2017 (ISO, 2017) was followed. Briefly, 40 ml of Tris (1 mol/l), glycine 0.05 M and beef extract 1% (TGBE) buffer was added to each sample. Samples were incubated at room temperature with constant rocking for 20 min. The pH of samples was monitored every 10 min and, if necessary, adjusted to 9.5 ± 0.5 with NaOH 10 N. The supernatant was transferred to a 50 ml tube and clarified by centrifugation at 10 000 g for 30 min at 5 °C. The supernatant was collected and adjusted to a pH of 7.0. A total of 0.25 volumes of 5× PEG/NaCl solution (500 g/ l PEG 8,000 (Sigma-Aldrich, USA), 1.5 mol/ l NaCl) were added to the recovered supernatant and homogenized by shaking for 60 s and incubated with constant rocking (approximately 60 oscillations min−1) at 5 °C for 60 min. The samples were centrifuged at 10,000 g for 30 min at 5 °C, the supernatant was discarded. The pellet was resuspended in 500 μl of PBS. Samples were subdivided for further RNase pre-treatment as capsid integrity assay.

### RNase pre-treatment

Prior to RNA extraction, the eluted virus (500 µl of PBS) was divided into a 100 µl sample that was discarded, a 200 µl no-RNAse control sample, and a 200 µl RNAse treated sample. *Sequence design (Primers and CRISPR crRNA)*.

All primers and crRNA were designed within the same region. All available sequences of the CW1 strain of MNV-1 at GenBank were aligned using Clustal Omega (Sievers et al., 2011). The complete sequenced genome from ATCC^®^ VR-1937™ was used as the main CW1 MNV-1 strain and a conserved region was determined through a multiple alignment. The polyprotein coding region of the RNA dependent RNA polymerase (RdRp) and the capsid protein was chosen as the qPCR and CRISPR target (ORF1 and ORF2). The qPCR primers and probe were designed using NCBI-Primer-BLAST and IDT-PrimerQuest™ Tool, along with quality analysis tools (Integrated DNA Technologies IDT-OligoAnalyzer™, Primer3). Isothermal Recombinase Polymerase Amplification (RPA) primers were designed following the TwistAmp^®^ Assay Design Manual (TwistDx™, Cambridge, England). Additionally, a T7 promoter was added to the 5’ end of the forward RPA primer for further RNA transcription. The crRNA for MNV-1 was designed following the published guidelines from original Cas13a-based detection developed (Kellner et al., 2019), and obtained from Integrated DNA Technologies IDT (IA, USA). The primers and crRNA are listed in **Table 1**.

**Table 1.**
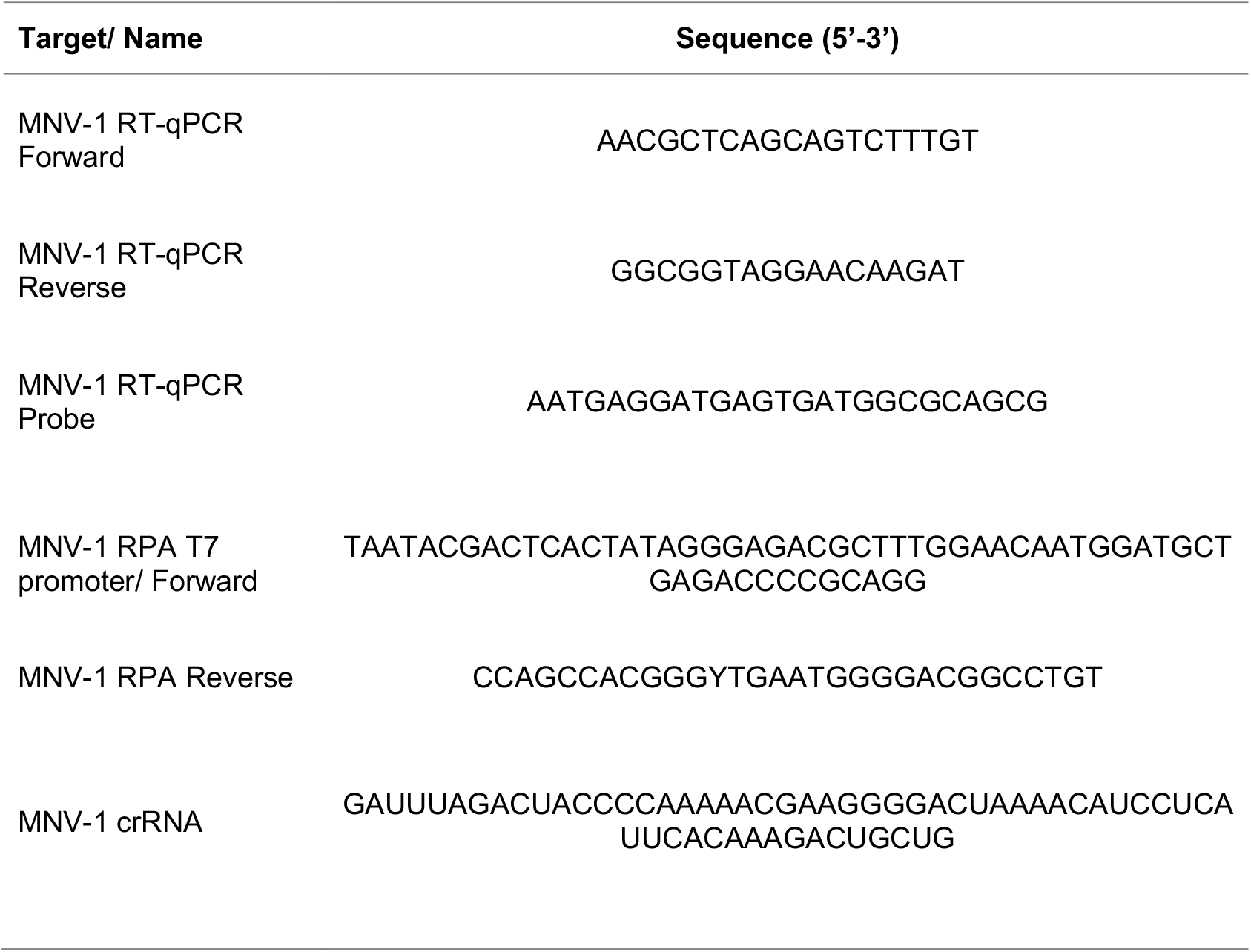
Sequences used in this study.

### Viral RNA extraction, reverse transcription, and qPCR

All viral RNA extractions were conducted on 140 µl of sample, post elution and RNAse/no-RNAse treatment, using the QIAamp viral RNA mini kit^®^ (Qiagen^©^, CA, USA), following the methodology described by the supplier. Viral RNA was eluted in 60 µl. The two step RT-qPCR was used for viral detection. Reverse transcription (cDNA) was carried out following supplier instructions from the High-Capacity cDNA Reverse Transcription Kit (Applied Biosystems™, CA, USA) at a final volume of 20 µl. The cDNA synthesis conditions were 10 min at 25 °C, 120 min at 37 °C and 5 min at 85 °C using a Veriti™ 96-Well thermal Cycler (Applied Biosystems™). Then the qPCR was performed using the SensiFAST™ Probe No-ROX Kit (Meridian Bioscience^®^, OH, USA) with final volume and concentration as recommended by the manufacturer. Cycling conditions at a Thermo Scientific™ PikoReal™ Real-Time PCR System (Thermo Scientific™, MA, US) were set at 3 min at 95 °C and 45 cycles of 10 s at 95 °C and 30 s at 60 °C; 3 µl of extracted viral RNA were used per reaction. For absolute quantification, a standard curve was constructed using a pure amplification product purified from an agarose gel, using the Wizard® SV Gel and PCR Clean-Up System (Promega^©^ WI, USA). Once purified, the DNA was measured by spectrophotometry using the NanoDrop™ 2000 (Thermo Scientific™). DNA were ten-fold diluted.

### Isothermal Recombinase Polymerase Amplification (RPA)

We followed the supplier instructions from the TwistAmp™ Liquid Basic Kit (TwistDx™, Cambridge, England), where in a final volume of 50 µl, 10 µM of each primer was added, plus 29.5 μl of primer free rehydration buffer, 5 µl of extracted viral RNA and, as reaction starter, 280mM of magnesium acetate (MgOAc). Samples were incubated at 39°C for 20 min and the amplification product was visualized on an agarose gel.

### Cas13a-based assay

CRISPR detection consisted of the combination of RPA amplification, T7 transcription, and Cas13a system into a single reaction, plus an RNase treatment prior to RNA extraction. A reaction volume of 50 μl was used with the following reagents and concentrations: 29.5 μl of RPA reaction (TwistAmp^®^ Liquid Basic kit, TwistDX, United Kingdom), 2.4 μl forward primer (10 μM), 2.4 μl reverse primer (10 μM), 0.25 μl of Cas13a (10μM) (SignalChem Biotech, Richmond, Canada), 0.125 μl crRNA (10 μM), 1.5 μl T7 RNA polymerase (50 U/μl; Thermo Scientific™, MA, US), 2 μl ribonucleoside triphosphate (rNTP) mix (50 mM; Thermo Scientific™, MA, US), 2 μl RNase inhibitor (40 U/μl; Promega© WI, USA), 0.5 μl MgCl2 (500 mM; Promega© WI, USA), 1 μl of RNaseAlert™ Lab Test Kit v2 (1 μM; Invitrogen™, MA, US), and RNase-free H_2_O. Then 3 μl of template and 2.5 μl of magnesium acetate (280 mM) were added to the tube cap and the tube cap closed. This mixture was vortexed and centrifuged to start the reaction. The fluorescence intensity was measured every min for 2 h in the PikoReal™ Real-Time PCR System (Thermo Scientific™, MA, US).

### Evaluation of the sensitivity and specificity of CRISPR detection

To evaluate the sensitivity of detection of our integrated method, six ten-fold serial dilutions ranging from 5.06 x 10^5^ to 5.06 x 10^0^ gc/ rx of MNV-1 were used to determine the limit of detection (LOD, CRISPR detection), and the limit of quantification (LOQ, RT-qPCR). The evaluate the specificity of our integrated method DNA, the amplification of MNR RNA was compared to the amplification from other non MNV-1 RNA targets including hepatitis A virus (HAV), human norovirus genogroups GI and II, and rotavirus. The HAV target was the 5′-UTR region for all HAV strains. Human norovirus genogroup GI and GII primers and targets were as described in the FDA BAM Chapter 26 (Williams-Woods et al., 2022). Both HAV and norovirus targets were generated using gBlocks gene fragment constructs (IDT, USA). The rotavirus target was the rotavirus region VP6 from rotavirus strain A using the pT7-VP6SA11 plasmid, a gift from Takeshi Kobayashi (Addgene plasmid # 89166; http://n2t.net/addgene:89166; RRID:Addgene_89166).

### Efficiency of recovery

For qPCR, the recovery efficiency of the elution and concentration steps was estimated using the equation: Virus recovery yields (%) = 10 ^(ΔCq/m)^ × 100% where ΔCq is the Cq value of extracted viral RNA from the food sample minus the Cq value of viral RNA extracted from the MNV inoculum dilution dose, and m is the slope of the virus RNA transcript standard curve. For the CRISPR method, the recovery efficiency was estimated using the equation: Virus recovery yields (%) = (mean fluorescence from eluted sample/ mean fluorescence from non-eluted sample) × 100 %. All assays were performed three separate times on three replicate samples. Statistical analyses were performed using the statistical packages in R programming language (R studio) using the t-test for independent samples where *p <* 0.05 was considered significant.

## 3. Results

### RT-qPCR of RNase pre-treated lettuce and blueberries samples

To quantify the limit of detection (LOD) of the RT-qPCR RNAse pretreatment MNV-1 detection assay, a ten-fold serial dilution of known concentrations were inoculated on lettuce (**Table 2**) and blueberry (**Table 3**) samples. The dilution range was 8.59 and 2.59 log10 genomic copies (gc) on 25 g of produce sample, equivalent to 5.06 × 10^6^ to 5.06 × 10^0^ gc/rx. The LOD for the RT-qPCR MNV-1 detection assay, without RNAse ONE™, for lettuce samples, was 2.59 log10 gc/25 g sample (5.06 gc/rx) inoculated MNV-1 (**Table 2**). At this dose, the average eluted MNV-1 was log_10_ 3.00 ± 2.44 gc/25 g. For blueberries samples, the LOD was tenfold higher at 3.59 log10 gc/25 g sample (5.06 × 10^1^ gc/rx) inoculated MNV-1 (**Table 3**). At this dose, the average eluted MNV-1 was log_10_ 2.65 ± 1.41 gc/ 25 g. At each dose, RNase treated samples, compared to untreated, had lower detectable RNA (**Table 2** and **3**). The RNase treated samples, compared to untreated samples, ranged in log reduction, across doses, between 76.72 ± 4.06 to 88.64 ± 7.21 % (lettuce) and 75.41 ± 6.25 to 83.08 ± 3.55 (blueberries). Heat inactivated MNV-1 at 80 °C was included as an positive RNase control. RNase treated, compared to untreated, MNV-1 at 80°C exhibited a reduction across doses (lettuce 93.5-95.3%; blueberries 96.1-96.8%).

**Table 2.** RT-qPCR detection of MNV-1 in inoculated lettuce samples.

| **Total Inoculum<sup>a</sup>** | **Lettuce** |  | **Lettuce RNase ONE™** |  |
| --- | --- | --- | --- | --- |
|  | **log10 gc/ 25g**<br>**± SD (Cq)** | **% Recovery**<br>**± SD<sup>b</sup>** | **log10 gc/ 25g**<br>**± SD (Cq)** | **% Reduction**<br>**± SD<sup>c</sup>** |
| 8.59 | 6.59 ± 6.29 (26.81) | 1.00 ± 0.50 | 5.95 ± 5.66 (28.72) | 76.72 ± 4.06 |
| 7.59 | 5.95 ± 5.67 (28.98) | 2.33 ± 1.21 | 5.01 ± 4.60 (31.85) | 88.64 ± 7.21 |
| 6.59 | 4.99 ± 4.74 (32.21) | 2.52 ± 1.39 | 4.25 ± 3.82 (34.47) | 81.92 ± 7.21 |
| 5.59 | 4.41 ± 4.28 (34.46) | 6.72 ± 4.89 | 3.48 ± 3.20 (37.83) | 88.25 ± 8.61 |
| 4.59 | 3.77 ± 3.57 (35.68) | 15.46 ± 9.44 | 2.93 ± 2.25 (39.44) | 85.87 ± 11.49 |
| 3.59 | 3.35 ± 3.29 (36.81) | 58.62 ± 50.56 | Not detected | Not detected |
| 2.59 | 3.00 ± 2.44 (38.68) | - | Not detected | Not detected |

<sup>a</sup> Total inoculum was calculated as MNV-1 genomic copies and expressed as log10 gc/ in 25 g.
<sup>b</sup> See methods. % recovery compared the MNV-1 gcs quantified after elution from the produce sample to the MNV-1 gcs used to inoculate the produce sample
<sup>c</sup> See methods. % reduction compared the MNV-1 gcs quantified after RNase treatment to the MNV-1 gcs quantified without RNase treatment. .

**Table 3.**
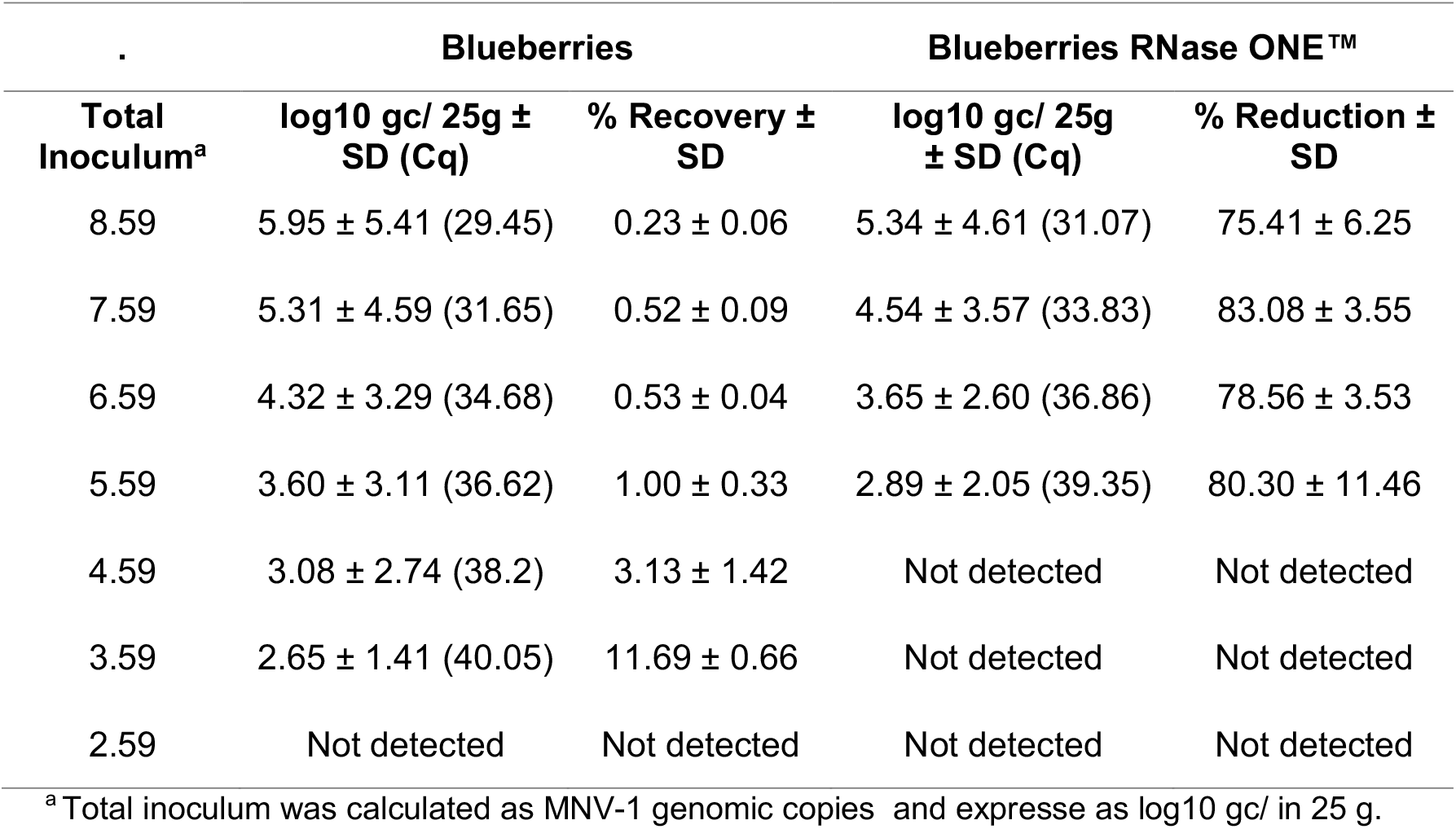
RT-qPCR detection of MNV-1 in inoculated blueberries samples.

| . | Blueberries |  | Blueberries RNase ONE™ |  |
| --- | --- | --- | --- | --- |
| Total Inoculum<sup>a</sup> | log10 gc/ 25g ± SD (Cq) | % Recovery ± SD | log10 gc/ 25g ± SD (Cq) | % Reduction ± SD |
| 8.59 | 5.95 ± 5.41 (29.45) | 0.23 ± 0.06 | 5.34 ± 4.61 (31.07) | 75.41 ± 6.25 |
| 7.59 | 5.31 ± 4.59 (31.65) | 0.52 ± 0.09 | 4.54 ± 3.57 (33.83) | 83.08 ± 3.55 |
| 6.59 | 4.32 ± 3.29 (34.68) | 0.53 ± 0.04 | 3.65 ± 2.60 (36.86) | 78.56 ± 3.53 |
| 5.59 | 3.60 ± 3.11 (36.62) | 1.00 ± 0.33 | 2.89 ± 2.05 (39.35) | 80.30 ± 11.46 |
| 4.59 | 3.08 ± 2.74 (38.2) | 3.13 ± 1.42 | Not detected | Not detected |
| 3.59 | 2.65 ± 1.41 (40.05) | 11.69 ± 0.66 | Not detected | Not detected |
| 2.59 | Not detected | Not detected | Not detected | Not detected |

<sup>a</sup> Total inoculum was calculated as MNV-1 genomic copies and expresse as log10 gc/ in 25 g.

### CRISPR Cas13a-based detection optimization; RPA detection and LOD (sensitivity)

The Cas13a-based detection is intended to be a one-pot detection that uses RPA isothermal pre-amplification in combination with Cas13a. We tested out RPA single reactions using viral RNA eluted from lettuce and blueberries samples. We found out that RPA isothermal amplification by itself is sensitive enough for the detection of MNV-1 at the lowest inoculated dilution of 5.06 × 10^0^ gc/rx in both, lettuce, and blueberries eluted samples. Before testing Cas13a-based detection in produce samples, we optimized the detection of MNV-1 along with crRNA concentration (e.g., 10 µM exhibited greatest signal yields), time of detection, limits of quantification (LOQ) and specificity with our target. We confirmed that MNV-1 was detected with all required reagents but not when any of the reagents were missing (**Figure 1A**). We selected 60 min as the cut-off time for detection because the fluorescence signal reached a plateau at that time. To quantify the LOQ of the Cas13a-based detection, ten-fold serially diluted MNV-1 (5.06 × 10^6^ to 5.06 × 10^0^ gc/rx) were tested (**Figure 1B**). The LOQ was at least 5 gc/ rx (total of 15 gc). Thus, a signal of 144.05 ± 107.26 fluorescence arbitrary units (a.u.) was the minimal positive signal of our method; samples below 107.26 fluorescence a.u. were considered negative. The specificity of the assay for MNV-1 was confirmed when the assay detected MNV-1 target but not synthetic DNA targets of GI and GII Norovirus, Hepatitis A virus and a plasmid rotavirus pT7-VP6SA11 (**Figure 1C**).

**Figure 1.**
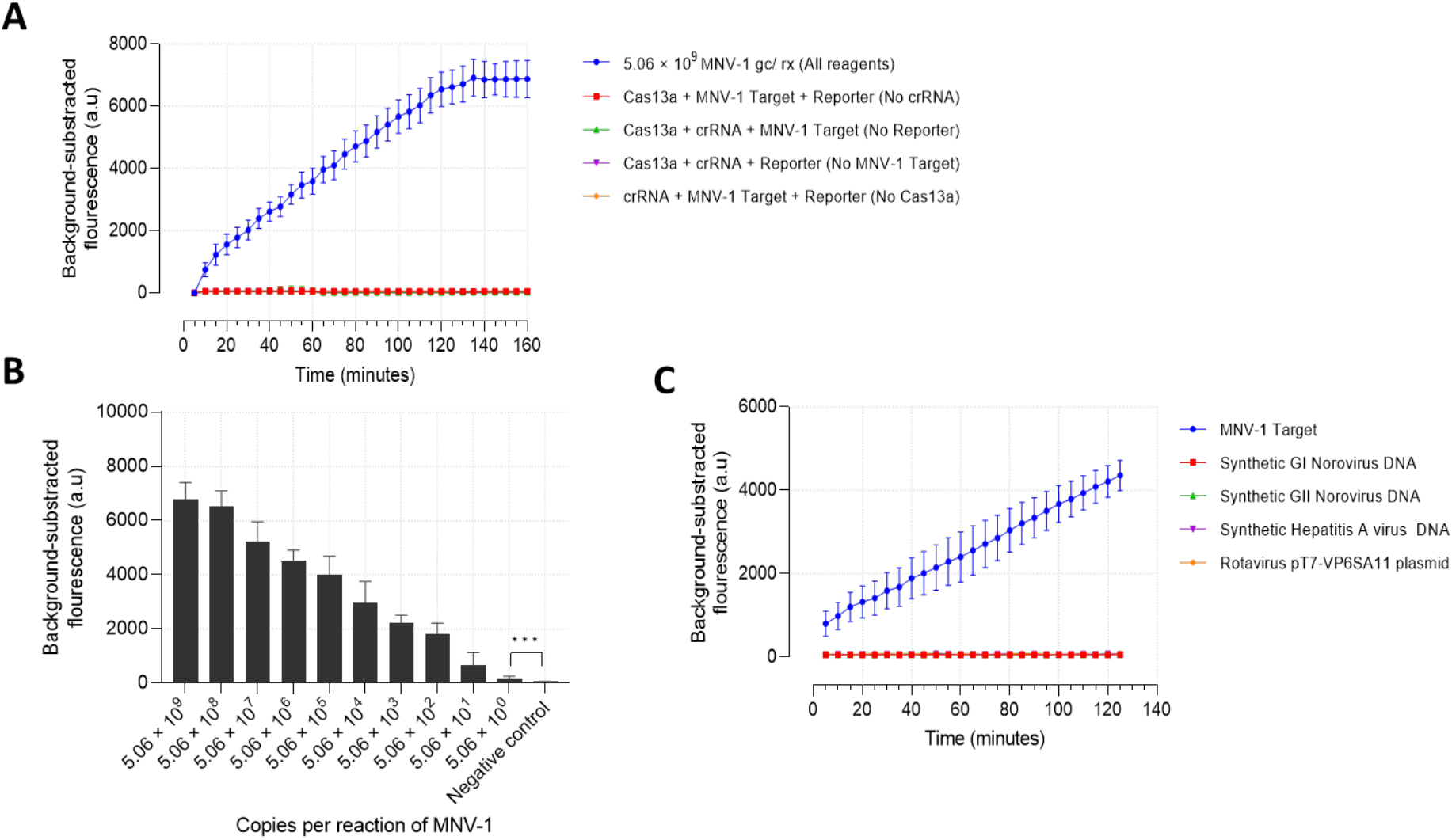
Cas13a based detection optimization. (**A**) The background-subtracted fluorescence signal increased over time and reached a plateau approximately at 120 minutes but only in a reaction mix with all required reagents. (**B**) The limit of detection was 5.06 for Cas 13a detection of the MNV-1 target.; *** *p <* 0.001. (**C**) Cas 13a based detection detected MNV-1 but not HAV, GI and GII norovirus synthetic targets and rotavirus VP6 containing plasmid (see Methods for details).

### CRISPR on lettuce and blueberries samples

The CRISPR method, consisting of the integration of RNase pre-treatment, RPA pre-amplification and Cas13a detection, was evaluated on inoculated lettuce (**Figure 2A**) and blueberry (**Figure 2B**) samples. Consistent with the prior RT-qPCR without RNase pre-treatment, our CRISPR-Cas13a method, without RNase pre-treatment, detected, for lettuce, 2.59 log10 gc/25 g sample (5.06 gc/rx) and, for blueberries, 3.59 log10 gc/25 g sample (5.06 × 10^1^ gc/rx) **(Table 4**). However, when the samples were pre-treated with RNase ONE™,we observed a positive signal corresponding to 3.59 log10 gc/25 g sample (5.06 × 10^1^ gc/rx) for both lettuce and blueberries, which was higher than the RT-qPCR assay (**Table 2** lettuce 4.59 log10 gc/25 g sample [5.06 × 10^2^ gc/ rx], **Table 3** blueberry 5.59 log10 gc/25 g sample [5.06 × 10^3^ gc/ rx]) (**Table 4**). RNase treated MNV-1 at 80°C (positive RNase control), compared to untreated, exhibited a reduction across doses for lettuce and blueberries..

**Table 4.** Comparison between the MNV-1 detection of RT-qPCR and CRISPR methods between lettuce and blueberries at each inoculated concentration.

| Inoculated concentration | RT-qPCR <sup>a</sup> |  |  |  | CRISPR <sup>a</sup> |  |  |  |
| --- | --- | --- | --- | --- | --- | --- | --- | --- |
|  | Lettuce |  | Blueberries |  | Lettuce |  | Blueberries |  |
|  | No RNase ONE™ | RNase ONE™ | No RNase ONE™ | RNase ONE™ | No RNase ONE™ | RNase ONE™ | No RNase ONE™ | RNase ONE™ |
| $5.06 \times 10^6$ | + | + | + | + | + | + | + | + |
| $5.06 \times 10^5$ | + | + | + | + | + | + | + | + |
| $5.06 \times 10^4$ | + | + | + | + | + | + | + | + |
| $5.06 \times 10^3$ | + | + | + | - | + | + | + | + |
| $5.06 \times 10^2$ | + | + | + | - | + | + | + | + |
| $5.06 \times 10^1$ | + | - | + | - | + | + | + | + |
| $5.06 \times 10^0$ | + | - | - | - | + | - | - | - |

<sup>a</sup> MNV-1 detected (“+”; Cq ≤ 40) and undetected (“-”, Cq > 40)

**Figure 2.**
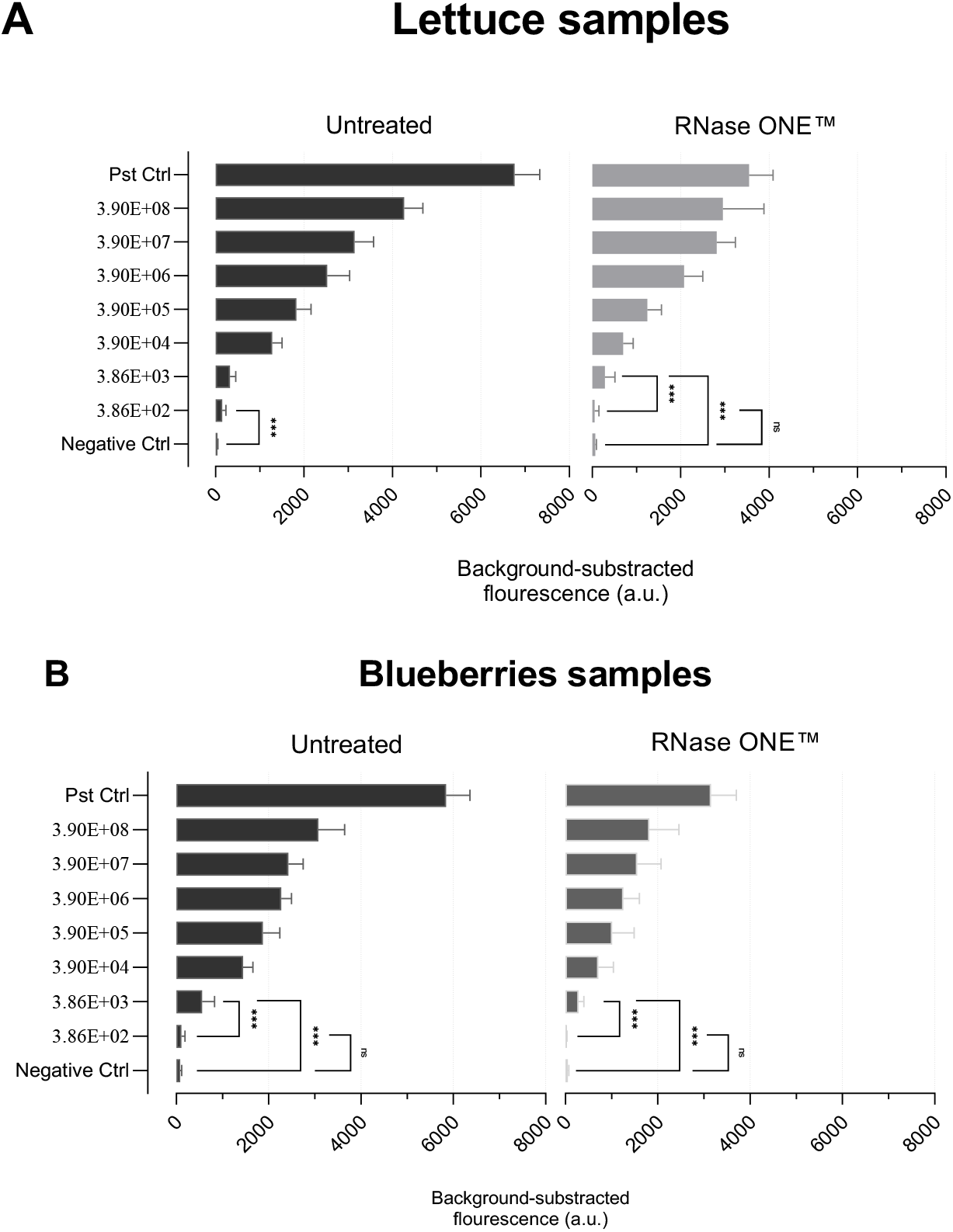
Cas13a based detection of inoculated MNV-1 on lettuce (**A**) and blueberry (**B**). In each panel, samples that were untreated were compared to samples that were RNAse treated. A signal of 144.05 ± 107.26 fluorescence arbitrary units (a.u.) was the minimal positive signal of our CRISPR method.

### Recovery efficiency

We compared the recovery efficiencies between lettuce and blueberries using RT-qPCR and CRISPR. The recovery yields range, across doses, of MNV-1 eluted and detected by RT-qPCR from lettuce samples was 1.0 to 58.6%, and from blueberries was 0.2 to 11.7% (**Table 2** and **3**). The recovery yields range, across doses of MNV-1 eluted and detected by CRISPR from lettuce samples was 1± 0.50 to 58.62 ± 50.56 %, and from blueberries was 0.23 ± 0.06 to 11.69 ± 0.66 %. By RT-qPCR, across doses, lettuce recovery yields were significantly higher (p < 0.05) than blueberry recovery yields. By CRISPR, across doses, there was no significant difference between lettuce and bluberry recovery yields.

## 4. Discussion

The goal of this study was to detect MNV-1 with an intact capsid, a proxy for infectivity, through a CRISPR-Cas13a based detection method in conjunction with a capsid integrity assay. We found that this CRISPR-based assay was both sensitive in detecting MNV-1 on lettuce and blueberries and did not cross-react with other genetic material. We also noted that this CRISPR-based, compared to the RT-qPCR-based, assay had the same produce-specific limit of detection (LOD) of MNV-1 on either lettuce (2.59 log10 genomic copies [gc]/ 25 g) or blueberry 3.59 log10 gc/ 25g) samples but had LOD differences by produce type. Lastly, CRISPR, compared to the RT-qPCR-based, assay after capsid integrity pre-treatment, detected MNV-1 RNA at a lower MNV-1 inoculation dose on lettuce and blueberry samples.

The CRISPR-based assay was both sensitive in detecting MNV-1 on lettuce and blueberries and did not cross-react with other RNAs from enteric viruses. The Cas13a/RPA isothermal amplification’s limit of detection (LOD) was 2.59 log10 gc/ 25g (5.06 gc/reaction, 1.69 gc/μl) in both lettuce and blueberries within 60 minutes. Other norovirus reports suggested higher or comparable LOD results employing either Cas13a or Cas12a detection and using fluorescence readers or lateral flow strips (LFS) for readout (Duan et al., 2022; Qian et al., 2021). For instance, Jia et al. (2020) reported an LOD of 50 norovirus GII gc/reaction in human stool samples within 20 minutes using RPA and a lateral flow test. Han et al. (2020) reported an LOD of 166 copies/μl GII norovirus in shellfish, water, and feces within 20 minutes using RT-RPA. Ma et al. (2018) reported an LOD of 100 gc of MNV-1 in mice fecal, fecal, and gastric tissue specimens within 16 minutes using RPA. Duan et al. (2022) reported an LOD of 5 gc/ reaction of GII.4 in stool samples within 40 minutes using RPA-Cas13a, a fluorescence reader, and LFS. Additionally, Li et al. (2024) reported an LOD of 2.5 gc/reaction of norovirus GII.4 and GII.17 in stool samples within 40 minutes using RPA-Cas13a and a portable blue light transilluminator. The lower detection time range (16-40 minutes) of these studies compared to our study’s detection time (60 minutes) could be attributed to the variation of fluorescence signal peaks (Ke et al., 2021) or the designed crRNA, which affects cleavage efficiency and fluorescence intensity (Gootenberg et al., 2017a; Ke et al., 2021; Leski et al., 2023; Tambe et al., 2018; Yang et al., 2023). The lack of cross-reaction of our assay with RNA from other enteric viruses was consistent with other CRISPR-based norovirus detection assays (Cas13a or Cas12a) for stool specimens, which also did not cross-react with other virus species such as rotavirus, enterovirus, and even other norovirus variants within the same GII genogroup (Duan et al., 2022; Qian et al., 2022).

This CRISPR-based and the RT-qPCR-based assay had the same produce-specific limit of detection (LOD) of MNV-1 on either lettuce (2.59 log10 gc/ 25g) or blueberry (3.59 log10/ 25g) samples but had LOD differences by produce type. The similarity in detection efficiencies between CRISPR- and RT-qPCR based assays, as we have observed, have been documented by other groups (Duan et al., 2022; Gootenberg et al., 2017). Both the CRISPR- and RT-qPCR-based assays exhibited a higher MNV-1 LOD on lettuce, over blueberry. We rejected the hypothesis that lettuce, over blueberry, samples had less detection inhibitors that similarly affected both the CRISPR-based and RT-qPCR-based assays. Evidence to reject this hypothesis includes the same LOD results, across CRISPR-based and RT-qPCR-based assays in lettuce and, separately, in blueberries (**Table 4**). Instead, we hypothesized that the MNV-1 detection was similar across assays, but that the MNV-1 viral elution was superior in lettuce, over blueberry, samples. In support of this hypothesis, we observed significantly higher overall MNV-1-specific percent recovery in lettuce, over blueberry samples (**Table 2, 3**). These findings suggest that the food type matrix characteristics may influence viral recovery. Similar to our results, one other group also observed superior extraction efficiencies of MNV-1, MS2, and Tulane Virus in lettuce over blueberries (Tang et al., 2023) .

CRISPR assay, compared to the RT-qPCR, assay could detect lower MNV-1 inoculated gcs of MNV-1 RNA from lettuce or blueberry samples that were eluted and treated with RNase. It is unlikely that CRISPR-based, over RT-qPCR-based, assays are more sensitive at detecting, after RNase treatment, intact RNA in intact MNV-1 capsids. The reason for this is that we have already shown and asserted, based on other’s evidence (Duan et al., 2022; Gootenberg et al., 2017), that both CRISPR-based and RT-qPCR-based assays have the same sensitivity at detecting MNV-1 RNA. Instead, we hypothesize that the addition of RNase resulted in incomplete reduction of free MNV-1 RNA and generation of MNV-1 RNA fragments. We define free RNA as RNase-accessible RNA outside a capsid or inside a damaged “permeable” capsid. In support of this hypothesis, we and others have shown that RNase treatment of inoculated MNV-1 on produce does not result in complete (100%) reduction of MNV-1 RNA. Our heat-treated 80°C controls, when comparing RNase treated to untreated samples, exhibited a 93.5-96.8% (1.19-1.86 Log_10_) reduction of MNV-1 RNA across MNV-1 doses on lettuce and blueberry samples. Marti et al. (2017) reported, in lettuce eluted samples using MNV-1, a 99.9998% (5.7 log_10_) reduction by RT-qPCR and using RNase A. Other studies in non-food samples have also reported variation 98.96-99.93% (0.03-0.45 log_10_) in the reduction of heat-inactivated noroviruses by using RNase pre-treatment (Li et al., 2012; Monteiro and Santos, 2018). This result suggests RNase did not destroy all accessible RNA or that heat treatment did not fully provide access of all RNA to RNase. If this hypothesis is shown to be valid, then both CRISPR-and RT-qPCR-based assays, combined with RNase treatment to detect intact capsids (i.e., infectious virus), may lead to false positive results: detection of incompletely degraded RNA fragments instead of protected RNA in intact capsids. To explain our results, we then hypothesize that CRISPR-based, over RT-qPCR-based, assays may be more effective at detecting RNase-generated MNV-1 fragments and thus more likely to generate false positive results (i.e., presumptive infectious virus). To address these hypotheses, our next areas of research are to develop methodologies to adjust RNase treatment results by the “effectiveness” of RNA reduction of free RNA (e.g., RNase treated controls) and improve RNase conditions and controls to eliminate, and confirm the elimination, of free RNA. We also aim to further validate these CRISPR-based results with MNV-1 infectivity results.

Our RNase reduction in MNV-1 inoculated lettuce and blueberries (75-89%, Log_10_ 0.57-0.91 reduction), measured by RT-qPCR, was higher than other published work. Monteiro and Santos (2018), using norovirus GII isolated from stools of infected patients with gastroenteritis and Hepatitis A virus (HAV) HM175/18f (VR-1402) from cell culture, achieved a mean of 0.03 log reduction in GII norovirus and 0.95 log reduction in HAV in non-food samples. Li et al. (2012), using MNV-1 from tissue culture, reported a log reduction of approximately 0.04 using MNV-1 suspension and RNase One RT-PCR *in vitro*. We hypothesize that either our, compared to others, RNase reaction conditions were superior at RNA reduction or that our eluted samples, compared to others, had more RNase-accessible RNA.

This study had strengths and limitations. The first strength of this study was that CRISPR based detection leverages the precision of Cas13a to achieve qualitative, sensitive, and specific detection of viral RNA, providing a reliable indicator of the presence of a target. The second strength of this study was the direct relationship between the concentration of target RNA and the detection capability of the enzyme (**Figure 1B**), laying the foundation for a dependable quantitative assay. The third strength was the evaluation of viral recovery and detection on different produce types across multiple doses. One limitation of this study was that we could not directly compare RT-qPCR results to CRISPR results (e.g., viral recovery) because it is currently qualitative and not yet capable of precise quantification without further optimization. A second limitation was the requirement for thorough calibration, including individual comparisons of slopes determined by linear regression models for varying target concentrations. A third limitation may the methodological inability to achieve 100% reduction of free RNA across sample matrices (e.g., different produce types), thus needing new methodologies to adjust for this confounder.

There are several next research areas to apply this assay to human norovirus on foods. One area is to enable quantitative analysis of pathogens in food samples by optimizing primer concentrations and reaction conditions to avoid saturation effects (e.g., in the pre-amplification step) and estimating the ratios of active Cas13 to the target RNA. A second area is to develop a calibration workflow that provides reliable parallel analyses of varying target concentrations. A third area is to validate the method across different food matrices to ensure its accuracy and reliability, including the quantification of viral elution efficiency for each food matrix. A long-term goal would be integrate CRISPR into the existing detection toolkit of food safety professionals in industry and government. This would include a cost-benefit analysis of its use versus traditional viral detection methods.

In conclusion, CRISPR Cas13a couple with a capsid integrity assay demonstrates a sensitive and specific detection of noroviruses on fresh produce. The ability of CRISPR to detect viruses is comparable to RT-qPCR without the need for expensive RT-qPCR equipment. We also demonstrate its potential of detecting RNA in intact capsids, a proxy for infectivity. This report provides a foundation for additional cost-efficient, rapid, and in-field detection assays for human foodborne viruses on foods to control the spread of foodborne outbreaks.

